# Disrupted Sphingosine-1-Phosphate Homeostasis Drives Nephrotoxicity in Sphingosine-1-Phosphate Lyase Insufficiency Syndrome (SPLIS)

**DOI:** 10.1101/2025.01.21.634100

**Authors:** Adam Majcher, Kathrin Buder, Ranjha Khan, Julie D. Saba, Thorsten Hornemann

## Abstract

Sphingosine-1-phosphate lyase insufficiency syndrome (SPLIS), also known as nephrotic syndrome type 14 (NPHS14), is an autosomal recessive disorder characterized by renal, neurological, dermatological, endocrine, and immunological symptoms. This condition is caused by loss-of-function mutations in the *SGPL1* gene, which encodes sphingosine-1-phosphate lyase (SGPL1p/SPL), the enzyme responsible for the terminal degradation of sphingosine-1-phosphate (S1P) in sphingolipid catabolism.

We investigated a novel case of SPLIS associated with a recently reported *SGPL1* mutation (*c*.*1084T>A; p*.*Ser362Thr*). Using stable isotope flux analyses, we demonstrated in patient-derived fibroblasts and *HEK293T* SGPL1 knockout models that SGPL1p deficiency does not consistently result in pathological S1P accumulation.

Instead, SPL-deficient cells are able to maintain steady-state S1P levels through two compensatory mechanisms:

Regulation of de novo sphingolipid synthesis via the ORMDL-ceramide axis.
Increased conversion of excess ceramides into glycosphingolipids.

However, when steady-state conditions are disrupted—either by external sphingolipid supplementation or by impairing homeostatic control—a pathological increase in intracellular S1P occurs in SPL-deficient cells.

In vivo, *Sgpl1*-/-mice exhibited significant urinary excretion of S1P and marked S1P enrichment in the kidneys. This pathological accumulation of S1P dysregulates cytoskeletal homeostasis, impairing renal epithelial formation. Based on these findings, we hypothesize that the reabsorption of urinary S1P contributes to toxic renal accumulation, providing an explanation for the nephrotoxicity observed in SPLIS and its association with nephrotic syndrome.

Importantly, we found that the cytoskeletal disruptions could be mitigated by inhibiting the Rho-ROCK signaling pathway using the clinically approved inhibitor Fasudil. These findings illuminate the pathophysiological basis of SPLIS nephrotoxicity and propose a promising pharmacological intervention strategy.

**Graphical abstract:** 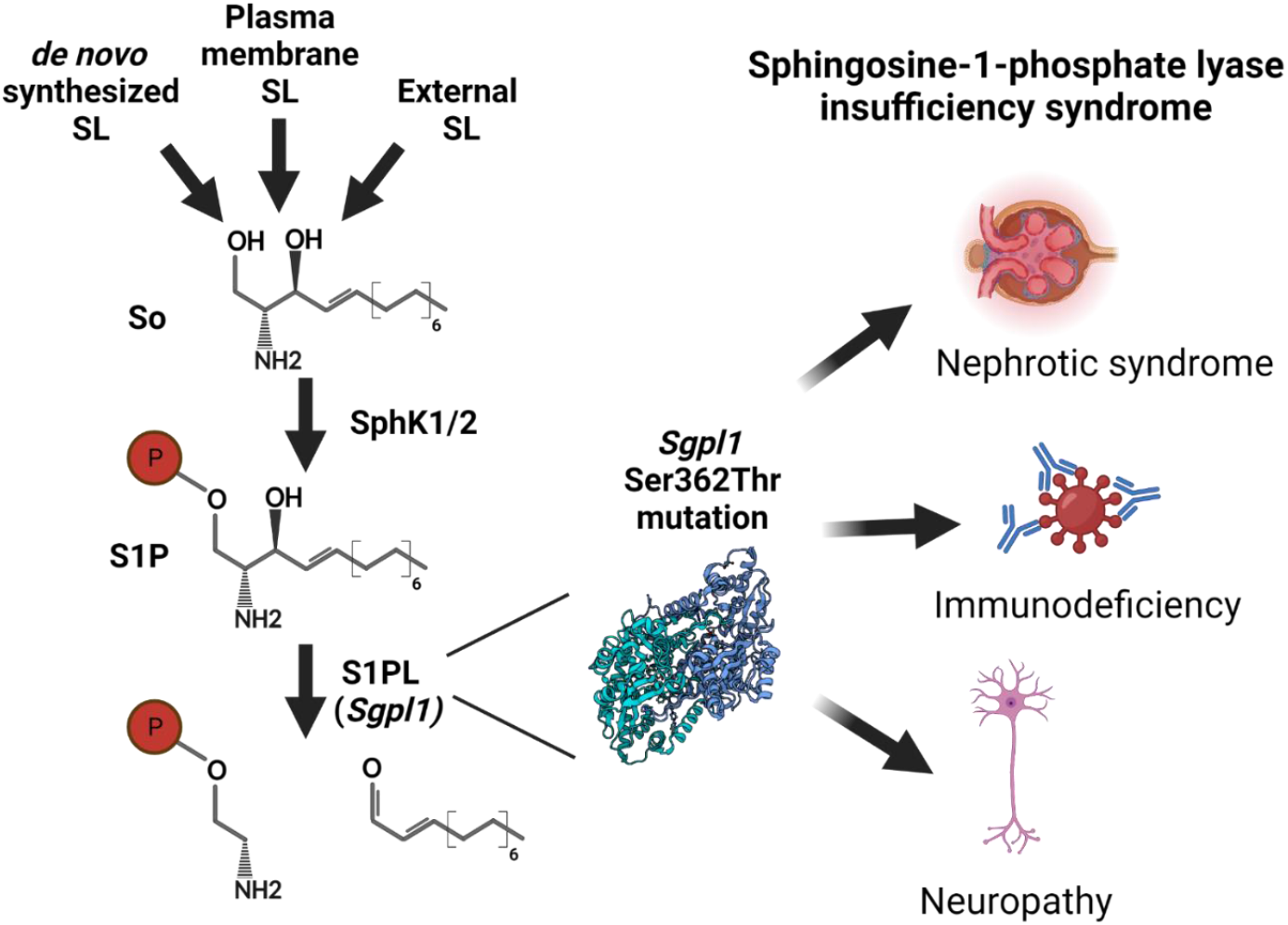

## Introduction

Sphingosine and its phosphorylated derivative sphingosine-1-phosphate (S1P) are critical bioactive lipids involved in numerous physiological and pathological processes.

Sphingosine-1-phosphate (S1P) is a potent lipid signaling molecule critical for regulating immune cell trafficking, vascular development, and maintaining endothelial integrity (1). S1P exerts its effects primarily through G protein-coupled receptors (S1PRs), which include five distinct subtypes (S1PR1–S1PR5), each with specific expression patterns and functions (1). In addition to its receptor-dependent actions, S1P also functions in a receptor-independent manner, influencing intracellular calcium homeostasis (2) and mitochondrial function (3). The terminal degradation of S1P is mediated by sphingosine-1-phosphate lyase (SGPL1p/SPL), which breaks it down into a long-chain aldehyde and phosphoethanolamine (4). SGPL1p is encoded by the *SGPL1* gene and is localized in the endoplasmic reticulum, where it catalyzes the final step in the catabolism of sphingolipids.

SPLIS or NPHS14 (also known as RENI syndrome), is a rare autosomal recessive disorder arising from biallelic mutations in *SGPL1* (5). Up to now, *SGPL1* mutations have been reported from over 76 patients (6). The disease is characterized by immunodeficiency as well as renal, neurological, skin and endocrine complications (7, 8). The overall mortality among SPLIS patients is 50%, primarily within the first decade of life, often due to end-stage kidney disease and sepsis (6).

Previous studies in cellular and mouse models have shown that SPLIS-associated SGPL1p variants have a reduced activity which results in increased S1P levels (9-11), suggesting that an aberrant S1P metabolism might contribute to the complications in SPLIS.

Opposing to SGPL1, the serine palmitoyl transferase (SPT) catalyzes the first and rate-limiting step of the SL *de novo* formation, opposes SPL activity. The SPT reaction typically conjugates L-Serine and Palmitoyl-CoA which forms the sphingoid base (keto)sphinganine, which is metabolized over several steps into Ceramides (Cer) and complex sphingolipids (4, 12). In addition to *de novo* synthesis as a source of sphingolipids, cells and tissues can also absorb free and phosphorylated sphingoid bases from extracellular sources. Cellular uptake of S1P and other phosphorylated sphingoid bases requires their dephosphorylation, catalyzed by phospholipid phosphatases (PLPP) followed by the passive absorption of the formed sphingosine (13). In addition, S1P can enter the cell via S1P receptor uptake. The resorbed sphingosine is then either reacylated to ceramide (salvage pathway) or again phosphorylated to S1P. Additionally, S1P is exported via sphingolipid transporter 2 (SPNS2), the major facilitator superfamily domain containing 2B (MFSD2B) (14). However, since *SGPL1* is ubiquitously expressed in all cells and tissues except erythrocytes and platelets, it is surprising that a systemic mutation in a universally active metabolic pathway manifests predominantly as a severe renal phenotype. In this study, we aimed to investigate why the reduction in SGPL1/SPL activity primarily affects kidney function. Specifically, we sought to identify the mechanisms that make kidney cells particularly vulnerable in SPLIS and to determine whether these underlying mechanisms can be targeted therapeutically.

## Methods

### Cell culture

HEK293T cell lines and primary fibroblasts were grown in high-glucose DMEM (Thermo Fisher Scientific) supplemented with 10% fetal bovine serum (FBS) and 1% Penicillin/Streptomycin (P/S) in a 5% CO_2_ incubator at 37°C. Primary skin fibroblasts were isolated from patient and healthy control skin biopsies. HK2 cell lines were grown in DMEM/F-12 (Thermo Fisher Scientific) supplemented with 10% fetal bovine serum (FBS) and 1% P/S in a 5% CO_2_ incubator at 37°C. For the fluorescence microscopy experiments, cells were grown in DMEM without phenol red supplemented with 10% fetal bovine serum (FBS) and 1% P/S in a 5% CO_2_ incubator at 37°C. For the transfection and silencing experiments, cells were grown in high-glucose DMEM supplemented with 10% fetal bovine serum (FBS) without any addition of P/S. Cells were tested for mycoplasma contamination.

### Generation of the *SGPL1* KO cell lines

HEK293T *SGPL1* KO cell line was a gift from Dr. Par Haberkant and generated using CRISPR/Cas 9 as described before (15).

Kidney proximal tubule cell line (HK2) *SGPL1* KO cell line was a gift from Dr. Robert Engel and generated using CRISPR/Cas9 as described before (16).

### Plasmids generation

Wild type *SGPL1* was cloned into the p.Lenti-V5 vector (V498–10, Thermo Fisher Scientific), and PCR-based site-directed mutagenesis was used to introduce SPLIS-associated *SGPL1* mutations using primers shown in Table 1, following manufacturer’s recommendation (Invitrogen). Generated plasmids were confirmed by Sanger sequencing.

**Table 1.**
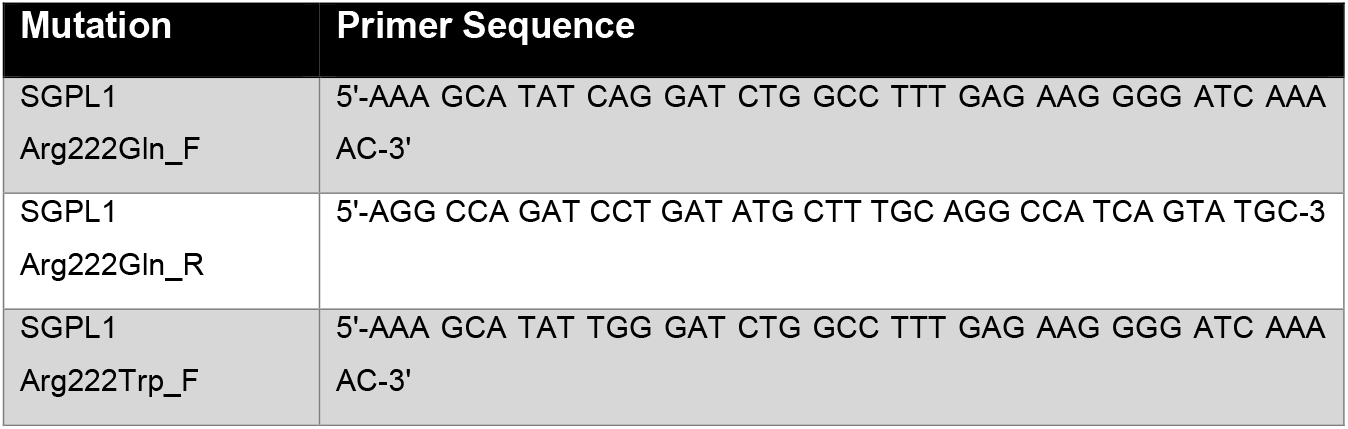
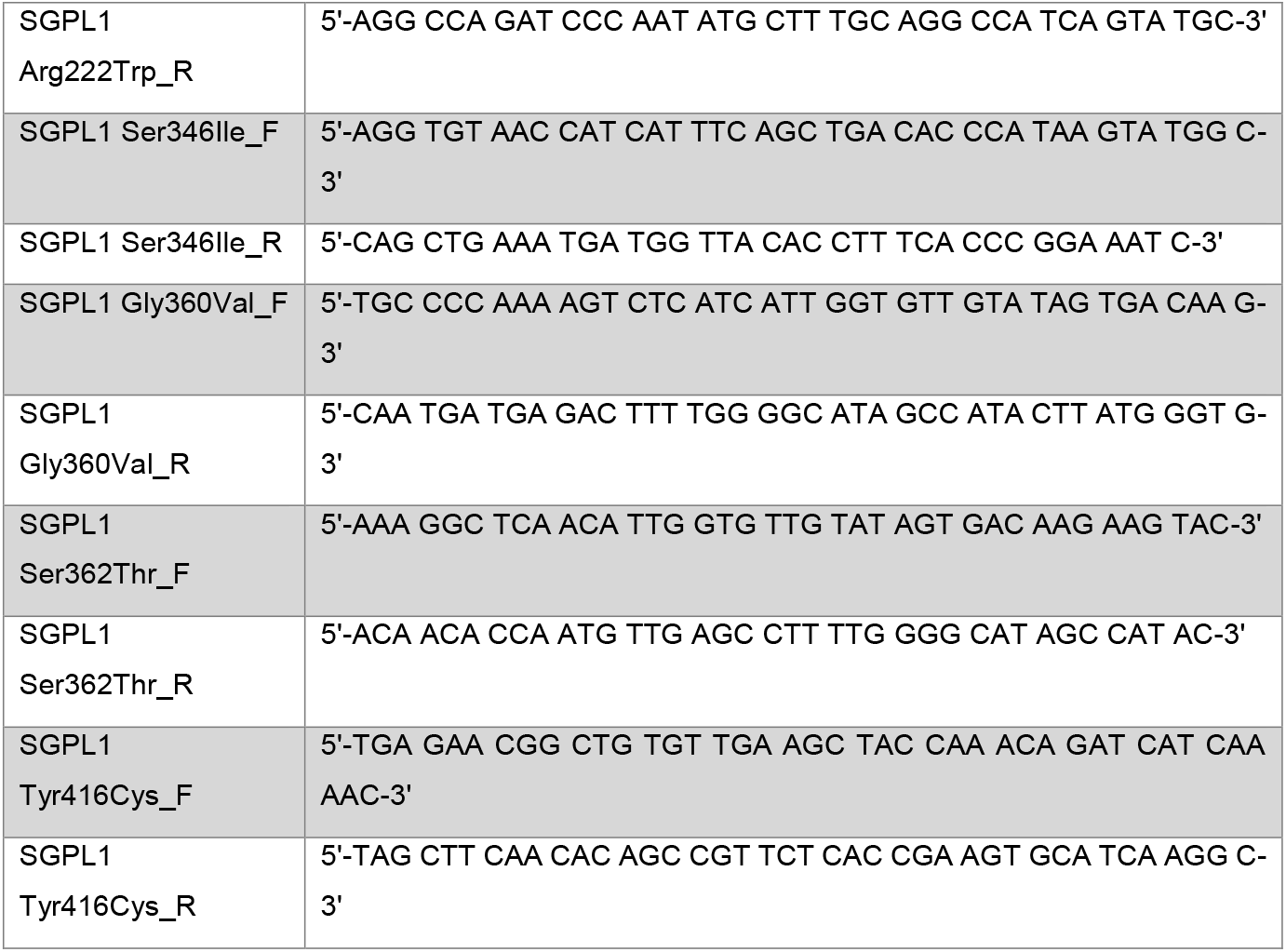
List of primers.

**Table 2.** Clinical phenotypes of the SPLIS (p.Ser362Thr) patient.

| Findings and symptoms |
| --- |
| **Kidneys:**<br>• Congenital nephrotic syndrome<br>• Anuric end-stage kidney disease* |
| **Immune system:**<br>• T-cell deficiency and reduced number of B cells<br>• Lymphopenia<br>• Hypogammaglobulinemia<br>• Recurrent viral and bacterial infections |
| **Central nervous system:**<br>• Sensorineural hearing loss<br>• Two generalized seizures at the age of 8 weeks and 8 months<br>• Computed tomography at the age of 8 months: slightly expanded cerebrospinal fluid spaces |
| • Global developmental delay |
| **Endocrinological system:**<br>• Adrenal insufficiency with recurrent episodes of arterial hypotension<br>• Hypothyroidism |
| **Lungs:**<br>• Recurrent respiratory tract infections with partial and global respiratory insufficiency<br>• Chest computed tomography at the age of 9 months: interstitial lung disease |
| **Electrolyte metabolism**<br>• Hypomagnesemia<br>• Hypocalcemia |
| **Additional findings**<br>• Poor feeding and failure to thrive<br>• Thrombophilia with catheter-associated thromboses<br>• Progressive decline in ultrafiltration in peritoneal dialysis |

### Generation of the *SGPL1* mutant cell lines

HEK293T *SGPL1* KO cell lines were transfected with P.Lenti-V5 Plasmid containing WT or mutant *SGPL1* using Lipofectamine 3000 (L3000001, Thermo Fisher Scientific). pLenti-V5 containing beta-Galactosidase (V498–10, Thermo Fisher Scientific) was used as an empty vector control. Transfected cells were selected by culturing in DMEM media (10% FBS) with Blasticidine (A1113903, Thermo Fisher Scientific, 20μg/mL) for 4 weeks.

### Silencing of ORMDL123

siRNAs targeting human *ORMDL1, 2, 3* were used to silence (knockdown) ORMDL1, 2, 3 expression. All siRNAs were mixed to an individual final concentration of 10 nM in reduced-serum media (Opti-MEM, 31985062, Thermo Fisher Scientific). Transfection was performed using Lipofectamine RNAiMAX transfection Reagent (13778150, Thermo Fisher Scientific) according to the manufacturer’s recommendations. The media was replaced after 24 hours with fresh DMEM (10% FBS), and cells were grown in total for 72 hours before the start of labeling experiments. Knockdown efficiency was determined using qRT-PCR.

### Isotopic labelling

For the SL labelling assay, cells were plated in 6-well plates. Cells were grown for 48 hours to 70% confluence in DMEM (10% FBS/1% P/S) (HEK293T, primary fibroblasts) or DMEM/F12 (10% FBS/1% P/S) growth media. 24 hours before harvesting, the medium was replaced with L-Serine free DMEM (C4331.0500, Genaxxon Bioscience) with 10% FBS/1% P/S and supplemented with 1mM of d_3_-^15^N-L-Serine (DNLM-6863-PK, Cambridge Isotope laboratories) with an addition of other treatments including (d7-Sa (860658), d7-So (860657) or d7-S1P (860659), Avanti Polar Lipids; FB1 (F1147), Sigma-Aldrich; Myr (M1177), Sigma-Aldrich) indicated in the figures. Cells were harvested by trypsinization, and cell pellets were washed 2 times with cold PBS. Cell pellets were then frozen and kept at -20°C until extraction.

### Mice tissues S1P measurement

Mice tissues were generated as described previously (17).

Tissue homogenates were prepared using Precellys homogenizer. Briefly, tissues were weighed on ice and transferred to the Precellys 2mL tubes with 6 beads per tube. MeOH was added to correct the concentration of the homogenates. Then, 9 rounds of homogenization at 5500 rpm, 25 seconds each round were performed at 4°C. Tissue homogenates were used for lipid extraction and analysis, as described below.

### Lipidomics analysis

Lipidomics analysis was performed as described previously (18). Briefly, extraction was performed by mixing (60 minutes, 37°C) the cell pellets, plasma, urine, or tissue homogenate with extraction buffer consisting of a mixture: methanol: methyl *tert-*butyl ether: chloroform 4:3:3 (v/v/v) and internal standards. After centrifugation (16 000 g, 10 min), the single-phase supernatant was collected, dried under N_2_, and stored at -20°C. Before analysis, lipids were dissolved in MeOH and separated on a C30 LC column using gradient elution with A) Acetonitrile: Water (6:4) with 10mM ammonium acetate and 0.1% formic acid and B) Isopropanol: Acetonitrile (9:1) with 10mM ammonium acetate and 0.1% formic acid at a flow rate of 260μL/minute. Eluted lipids were analyzed on a Q-Exactive HRMS (Thermo Fisher Scientific) in positive and negative modes using heated electrospray ionisation (HESI). MS2 fragmentation spectra were recorded in data-dependent acquisition mode. Peak integration was performed with TraceFinder 4.1 (Thermo Fisher Scientific). Lipids were identified by predicted mass (resolution 5ppm), retention time, and specific fragmentation patterns. Next, lipid concentrations were normalized to the internal standards (one per class) and total protein amount.

List of internal standards:

d5-1-deoxymethylsphinganine (SPB17:0;O, 860476, Avanti Polar Lipids) 100pmol/sample

C17 Sphingosine-1-phosphate (SPBP17:0;O2, LM2144, Avanti Polar Lipids) 50pmol/sample

1-deoxydihydroceramide (Cer18:0;O/12:0, 860460P. Avanti Polar Lipids) 100pmol/sample

1-deoxyceramide (Cer18:1;O/12:0, 860455, Avanti Polar Lipids) 100pmol/sample Dihydroceramide (Cer18:0;O2/12:0, 860635, Avanti Polar Lipids) 100pmol/sample Ceramide (Cer18:1;O2/12:0, 860512, Avanti Polar Lipids) 100pmol/sample

SM (SM18:1;O2/12:0, 860583, Avanti Polar Lipids) 100pmol/sample

Glucosylceramide (GlcCer18:1;O2/ /8:0, 860540, Avanti Polar Lipids) 100pmol/sample

SPLASH standard (330707, Avanti Polar Lipids) 2,5μL/sample

Transitions used for the identification of Sphingolipids:

Ceramides:

[M+H]^+^ → [M+H - H_2_O]^+^, [M+H]^+^ → [M+H - H_2_O - FA]^+^, [M+H]^+^ → [M+H - 2xH_2_O - FA]^+^

HexosylCeramides:

[M+H]^+^ → [M+H - Hexosyl]^+^, [M+H]^+^ → [M+H - H_2_O - FA - Hexosyl]^+^, [M+H]^+^ → [M+H

- 2xH_2_O - FA - Hexosyl]^+^

Sphingomyelins

[M+H]^+^ → [PO4-Choline]^+^, [M+H]^+^ → [M+H - H_2_O - FA – PO4-Choline]^+^, [M+H]^+^ → [M+H - 2xH_2_O - FA - PO4-Choline]^+^

Sphingosine-1-phosphates

[M+H]^+^ → [M+H - 2xH_2_O - FA – PO4]^+^, [M+H]^+^ → [M+H - H_2_O - FA – PO4]^+^

Free long-chain bases (Sphingosine and Sphinganine):

[M+H]^+^ → [M+H - H_2_O]^+^, [M+H]^+^ → [M+H – 2xH_2_O]^+^

Phosphatidyl cholines

[M+H]^+^ → [PO4-Choline]^+^, [M+H]^+^ → [Choline]^+^ [M+Formate]^+^ → [M+Formate - PO4-Choline - FA]^+^ Note: FA represents corresponding fatty acyl.

### Protein determination

For the protein content normalization, cell pellets remaining after the lipid extraction were dissolved in urea (8M) containing 1% 2-mercaptoethanol and mixed at 800 rpm at 90°C (Thermomixer (Eppendorf)) for 20 minutes. Next, protein extracts were snap frozen using dry ice, and the whole process was repeated 3 times. After centrifugation (16 100 rpm, 10 minutes), supernatants were collected and used for further analysis.

Total protein amounts were determined using Bradford assay (Bio-Rad, 1:50 dilution) according to the manufacturers’ recommendation.

### Western blotting

Protein isolation and Western blot were performed as described elsewhere (18). Primary antibodies used for immunodetection of V5-antigen were V5 Tag antibody (R960-25, Thermo Fisher Scientific, 1:2000 dilution, 1 hour at room temperature) and for Calnexin immunodetection, Anti-Calnexin (ZRB1147, Sigma-Aldrich, 1:2500 dilution, 1 hour at room temperature). As secondary antibodies horseradish peroxidase–conjugated anti-mouse (NC2294470, Thermo Fisher Scientific; 1:10 000, 1 hour at room temperature) and anti-rabbit (12-348, Thermo Fisher Scientific; 1:10 000, 1 hour at room temperature) IgGs were used.

### Toxicity assays

For the toxicity assay cells were grown for 72 hours in the 96-well plates with treatment (So (860490), Avanti Polar Lipids; FB1 (F1147), Sigma-Aldrich; Genz (5.38285), Sigma-Aldrich), as indicated in the figures. Next CellTiter-Glo **®** Luminescent Cell Viability Assay (G7570, Promega) was performed according to the manufacturer’s recommendations. Chemiluminesce signal was detected using TECAN infinite M 200 Pro reader. All conditions were corrected for the solvent concentration.

### Live cell microscopy

For the live cell microscopy, cells were grown in 96-well plates in the incubator at 5% CO_2_ and 37°C. Indicated treatment (JTE 013 (J4080), Fasudil (CDS021620), FTY720 (SML0700), Sigma-Aldrich) was added by media exchange. Then, the plate was immediately transferred to a live-cell imaging microscope (Olympus IX81) with a motorized stage, fitted with an incubator with pre-heated humidified atmosphere (Ibidi mixer), and kept at 37°C and 5% CO_2_. Phase-contrast images were acquired at 20x magnification every 15 minutes for 48 hours. Four images per well were acquired and compiled. Percentage contracted cells was acquired by selecting four 10mm rectangles (regions of interest) per well and then contracted cells and non-contracted cells were counted manually using ImageJ software. The same positions of the rectangles per well were used at all timepoints and conditions. Percentage of contracted cells per well was calculated as average of four percentages in rectangles. All original images of cut rectangles used for manual counting are available.

### Scratch assay

Scratch assay was performed as described previously (19). Briefly, fibroblasts were seeded in 12-well plates for 48 hours. Next, cell proliferation was stopped by 2-hour incubation with Mitomycin C (M4287, Sigma-Aldrich, 10 μg/ml). Then, a scratch was introduced in the middle of the well by a 20 μL pipette tip. In order to remove the cell debris, wells were rinsed once with PBS and fresh DMEM (10% FBS / 1% P/S) supplemented with given treatment was added. Then, plates were transferred to the live-cell imaging microscope (Olympus IX81) with a motorized stage, fitted with an incubator with pre-heated humidified atmosphere (Ibidi mixer, Ibidi GmbH), and kept at 37°C and 5% CO_2_. Phase-contrast pictures were acquired at 10x magnification every 30 minutes for 48 hours.

### Fluorescence microscopy

For the fluorescence microscopy cells were grown in 96-well plates in the incubator at 5% CO_2_ at 37°C. Indicated treatments were added by exchange of the media. After the given time (indicated in the figures), cells were washed 3 times with PBS and fixed using 4% PFA for 30 minutes. Next, cells were washed 3 times with PBS and then incubated for 1 hour with DAPI (D9542, Sigma-Aldrich, 1 μM in PBS) and Phalloidin-665 (18846, Sigma-Aldrich, 0.5 μM in PBS). Further, cells were washed with PBS and imaged immediately. Images were acquired using a fluorescence microscope (Olympus IX81) with a motorized stage at the 20x magnification.

### Periodic acid-Schiff staining (PAS)

PAS staining was performed on 5 micron kidney sections from *Sgpl1*^*-/-*^ and WT mice as we described previously (Khan et al IJMS). A total of 50 glomeruli and surrounding tissues were evaluated to quantify abnormalities including segmentally sclerosed glomeruli, glomeruli displaying mesangial hypercellularity, and interstitial fibrosis.

### Image analysis

Image analysis was performed using CellProfiler 4.2.1 using an in house made pipeline. First, the Nucleus area was segmented as a primary object (Otsu thresholding method) using DAPI channel image. Second, cellular area was identified and segmented as a secondary object from Nuclear area in Phalloidin channel image (Propagation method using Minimum Cross-Entropy thresholding). Then, log10(Cellular area/Nucleus area) was calculated per object and the average per well was used for further quantification.

### Data analysis, statistics and figures

All the data analysis and figure preparations were performed using GraphPad Prism 9.5.1 (Bar graphs, Stack plots) and Excel. Statistical analysis was performed in GraphPad Prism 9.5.1. The individual statistical methods used are indicated in the figures. Illustrations were made using BioRender webpage (BioRender.com). Final figures were prepared using Affinity Designer 2.

## Results

### A novel SPLIS-associated *SGPL1* p.(Ser362Thr) variant

An 8-week-old girl, term-firstborn of healthy Iraqi consanguineous parents (oligohydramnios, small for gestational age with birth weight of 2140 g at 37+1 gestational weeks), was diagnosed with dialysis-dependent end-stage kidney disease due to congenital nephrotic syndrome presenting with heavy proteinuria, hypalbuminemia and edema. Kidney biopsy and steroid treatment were not performed, considering the patient’s young age which was suggestive for a genetic cause of the nephrotic syndrome.. Whole exome sequencing identified a homozygous, variant in exon 12 of the *SGPL1* gene (c.1084T>A, p.Ser362Th) suspected to be pathogenic and causing SPLIS (Figure 1 B, C, D) (6). In the protein sequence, the mutation is located close to the cofactor-binding lysine at K353. Besides the renal phenotype, the patient revealed multiple extra renal manifestations summarized in Table 1. Adrenal calcifications identified initially, are shown in Figure 1. The clinical course was complicated by prolonged hospitalizations due to recurrent respiratory tract infections with global respiratory insufficiency, recurrent cytomegalovirus viremia and coagulase-negative staphylococci bacteremia, life-threatening arterial hypotension, catheter-related thromboses of several central veins of the upper thoracic aperture, increased need of peritoneal dialysis due to ultrafiltration insufficiency. With 8 months, after clinical presentation of a generalized seizure, cerebral computer tomography showed slightly expanded cerebrospinal fluid spaces. The patient died at the age of 9 months most likely due to a septic shock with subsequent multiorgan failure.

**Figure 1.**
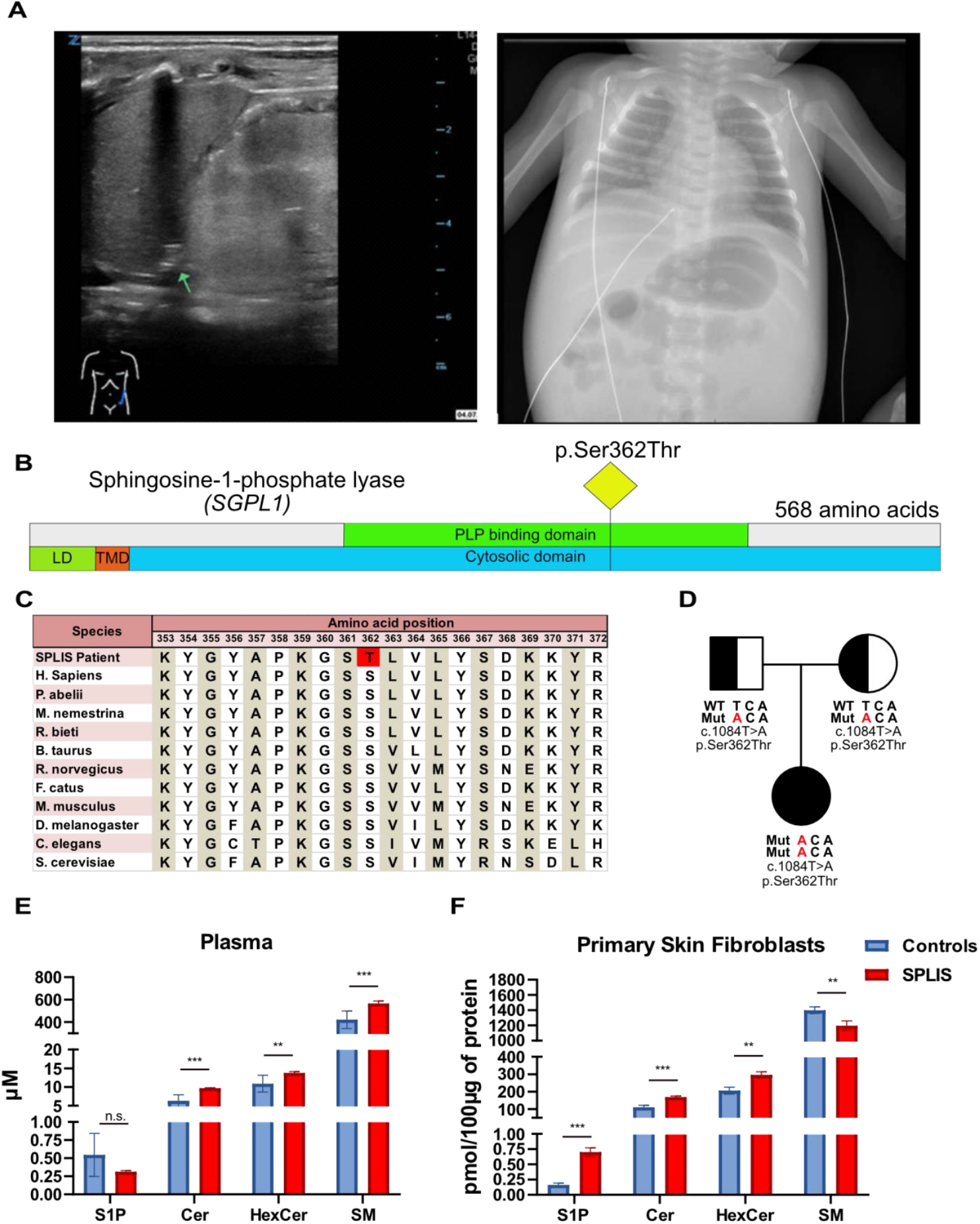
Clinical and molecular characterization of the new SPLIS-causing SGPL1 mutation p.Ser362Thr. **(A)** Sonographic and X-ray imaging of a patient with the *SGPL1* mutation (p.Ser362Thr). Left: Sonographic image showing a hyperechogenic kidney and adrenal calcifications (arrow). Only the left kidney and adrenal gland are depicted. Right: X-ray image revealing adrenal calcifications. **(B)** Schematic representation of the human *SGPL1* gene with an up-to-scale domain architecture. The SPLIS-associated mutation is marked with a yellow rectangle. **(C)** Conservation analysis of amino acid residues surrounding the SPLIS-causing mutation in *SGPL1*. The patient mutation (p.Ser362Thr) is highlighted in red. **(D)** Pedigree of the affected family, illustrating the segregation of the *SGPL1* variant (*NM_003901*.*3* c.1084T>A, p.Ser362Thr, Chr10(hg19):g.72633132T>A). **(E)** Plasma sphingolipid (SL) profile of the SPLIS (p.Ser362Thr) patient (N=3, technical replicates) compared to healthy controls (N=7). **(F)** SL profile of primary skin fibroblasts from the SPLIS patient versus healthy controls (N=3). Bar plots represent the mean ± SD of sums for each SL class. Statistical significance was assessed using a t-test, with multiple testing correction applied via the two-stage step-up method (Benjamini, Krieger, and Yekutieli). n.s. = non-significant; * = q < 0.05; ** = q < 0.005; *** = q < 0.0005.

To investigate whether the new *SGPL1* variant was associated with changes in overall plasma SL profile, we performed an untargeted LC-MS/MS based lipidomics analysis of patient’s plasma compared to healthy controls (N=7). In our analysis, we did not see significantly elevated S1P levels while Cer, HexCer and SM levels were slightly increased in the patient plasma. (Figure 1E). However, due to the patient’s very young age and severe condition, only a very limited amount of plasma was available. As a result, S1P levels were quantified from the general lipidomics analysis, which is not the best suited method to measure phosphorylated sphingoid bases. Unfortunately, there was insufficient plasma to verify S1P levels through a separate more sensitive, targeted S1P analysis. Patient derived skin fibroblasts showed a significant increase in S1P, Cer and HexCer while total SM was decreased (Figure 1F). However, the absolute increase in S1P was small (0.005pmol/μg protein) compared to the total differences seen for HexCer (1pmol/μg protein) or SM (2pmol/μg of protein). A similar change in the SL profile was also seen in HEK293T-*SGPL1 KO* cells (data not shown).

### Downregulation of SL de-novo synthesis prevents S1P accumulation in SGPL1p/SPL-deficient cells

SPT activity and de novo sphingolipid (SL) synthesis are regulated by the subunits ORMDL1, ORMDL2, and ORMDL3 in response to intracellular ceramide levels (20). To see whether SL *de novo* synthesis was altered in SGPL1p/SPL-deficient cells, we performed a stable isotope-labeling assay using d_3_-^15^N-Serine. Primary SPLIS fibroblasts as well as HEK293T *SGPL1* KOs cells showed a decreased SL *de novo* synthesis compared to controls. Supplementing cells with exogenous d7-sphingosine (d7-So) further reduced de novo sphingolipid (SL) synthesis in both KO and WT cells, indicating that an increase in intracellular SL levels leads to an enhanced compensatory downregulation of de novo synthesis (Figure 2B,C). After silencing ORMDL1-3 expression by siRNA we observed an increased *de novo* formation and a substantial accumulation of Sa1P and S1P (here shown as sum of both) in SGPL1p/SPL-deficient cells (Figure 2D, Supplementary Figure 1).

**Figure 2.**
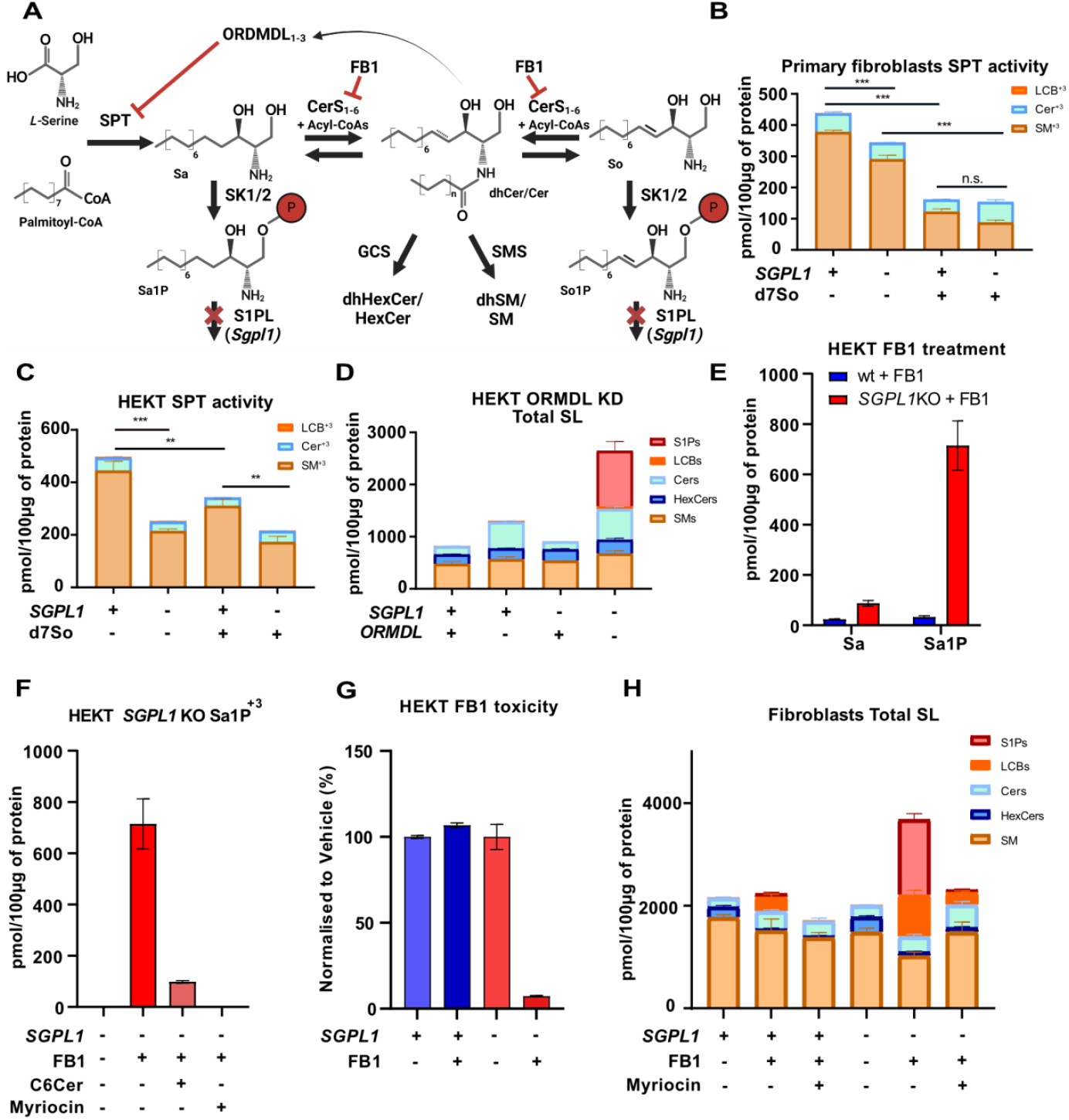
SPT regulation via the ORMDL-Cer axis prevents toxic S1P accumulation in SGPL1-deficient cells. **(A)** Schematic illustration of de novo sphingolipid (SL) synthesis regulation by serine palmitoyltransferase (SPT). **(B)** Reduced SPT activity (de novo synthesis) in SPLIS primary skin fibroblasts (SGPL1 -) compared to control fibroblasts (SGPL1 +). Cells were also supplemented with external SL (+) or vehicle (-) to assess homeostatic control. **(C)** SPT activity in *HEK293T* SGPL1 KO cells (-) compared to WT cells (+), with additional supplementation of d7-sphingosine (d7-So). **(D)** Total SL levels following ORMDL1-3 knockdown in *HEK293T* SGPL1 KO and WT cells. Cells were transfected with Lipofectamine 3000 using either scrambled (ORMDL +) or ORMDL-targeting siRNAs (ORMDL -) for 72 hours before initiating the experiment. **(E)** Total sphinganine (Sa) and sphinganine-1-phosphate (Sa-1-P) levels in the presence of fumonisin B1 (FB1) in *HEK293T* SGPL1 KO and WT cells. **(F)** De novo formation of Sa-1-P in *HEK293T* SGPL1 KO cells treated with vehicle, FB1, FB1 + cell-permeable C6-ceramide (C6Cer), or FB1 + SPT inhibitor Myriocin. **(G)** FB1 toxicity assay in *HEK293T* SGPL1 KO and WT cells. Cells were treated with FB1 for 72 hours, and total ATP levels were quantified using the CellTiter-Glo assay. **(H)** Total SL levels in SPLIS primary skin fibroblasts (SGPL1 -) or control fibroblasts (SGPL1 +) after treatment with vehicle (MeOH, FB1 -), FB1 (FB1 +), or FB1 + Myriocin (Myriocin +). SPT activity was quantified by measuring the incorporation of d3-15N-serine after 24 hours. Bar and stack plots represent all SL class species as mean ± SD (N=3). Hexosylceramides (HexCer) represent the sum of galactosylceramide and glucosylceramide. SL levels were analyzed using LC-MS/MS following lipid extraction. Statistical significance was determined using a Student’s *t*-test, with multiple testing correction applied via the two-stage step-up method (Benjamini, Krieger, and Yekutieli). * = p < 0.05; ** = p < 0.01; *** = p < 0.001. Data from the toxicity assay (**G**) were normalized to the average of vehicle-treated cells (N=4).

In a second approach, we modulated the homeostatic control by adding Fumonisin B1 (FB1), a pan-inhibitor of Ceramide Synthases 1-6 (CerS1-6) that prevents the *de novo* formation of dihydroceramide and subsequently ceramide downstream of SPT. FB1 leads to a substantial accumulation of *de novo*-formed Sa and Sa1P in *SGPL1* KO but not in WT cells (Figure 2E). This accumulation was significantly reduced when activating the inhibitory SPT subunits ORMDL1-3 with cell-permeable short-chain C6-Ceramide or by inhibiting SPT with Myriocin (Figure 2F). In comparison, FB1 was significantly more toxic for *SGPL1* deficient cells compared to WT (Figure 2G). In presence of FB1, phosphorylated and non-phosphorylated LCBs increased only in SGPL1p deficient cells (Figure 2H).

### Synthesis of higher SL is an “escape” mechanism for preventing toxic S1P accumulation in SPLIS

o circumvent SPT regulation by the ORMDLs, we added isotope-labeled d7-sphinganine (d7-Sa) directly to the cells and monitored the isotopic flux. The downstream products formed, such as d7-ceramide (d7-Cer), d7-hexosylceramide (d7-HexCer), and d7-sphingomyelin (d7-SM), showed a 3–4-fold increase (Figure 3A). Additionally, supplementation with d7-sphingosine (d7-So) or d7-sphingosine-1-phosphate (d7-S1P) at concentrations of 0.5 μM or 2.5 μM resulted in a significant increase in d7-Cer, d7-HexCer, d7-SM, and d7-S1P levels (Figure 3B). Total sphingolipid (SL) levels were calculated as the sum of all non-labeled and M+7-labeled SL species. Overall, SPLIS primary fibroblasts, *HEK293T* SGPL1 KO cells, and HK2 SGPL1 KO cells exhibited elevated SL levels compared to WT cells. Notably, in addition to S1P, ceramide (Cer) and hexosylceramide (HexCer) levels were predominantly increased, while total sphingomyelin (SM) levels showed minimal changes (Figures 3D and 3E). Total S1P levels were elevated in all three KO cell lines. However, all KO cell lines accumulated substantial amounts of HexCer (depicted in blue in Figures 3C, D, and E). To assess whether the conversion of excess SLs to HexCer is crucial for maintaining SL homeostasis and avoiding toxic SL accumulation in SPLIS, we incubated WT and SGPL1 KO cells with increasing concentrations of sphingosine (So) in the presence of a glucosylceramide synthase (GCS) inhibitor (Genz-123346). Blocking GCS indeed sensitized SGPL1 KO cells to So toxicity, indicating that the conversion of excess SLs into HexCer acts as a metabolic escape mechanism to prevent toxic S1P accumulation (Figure 3F).

**Figure 3.**
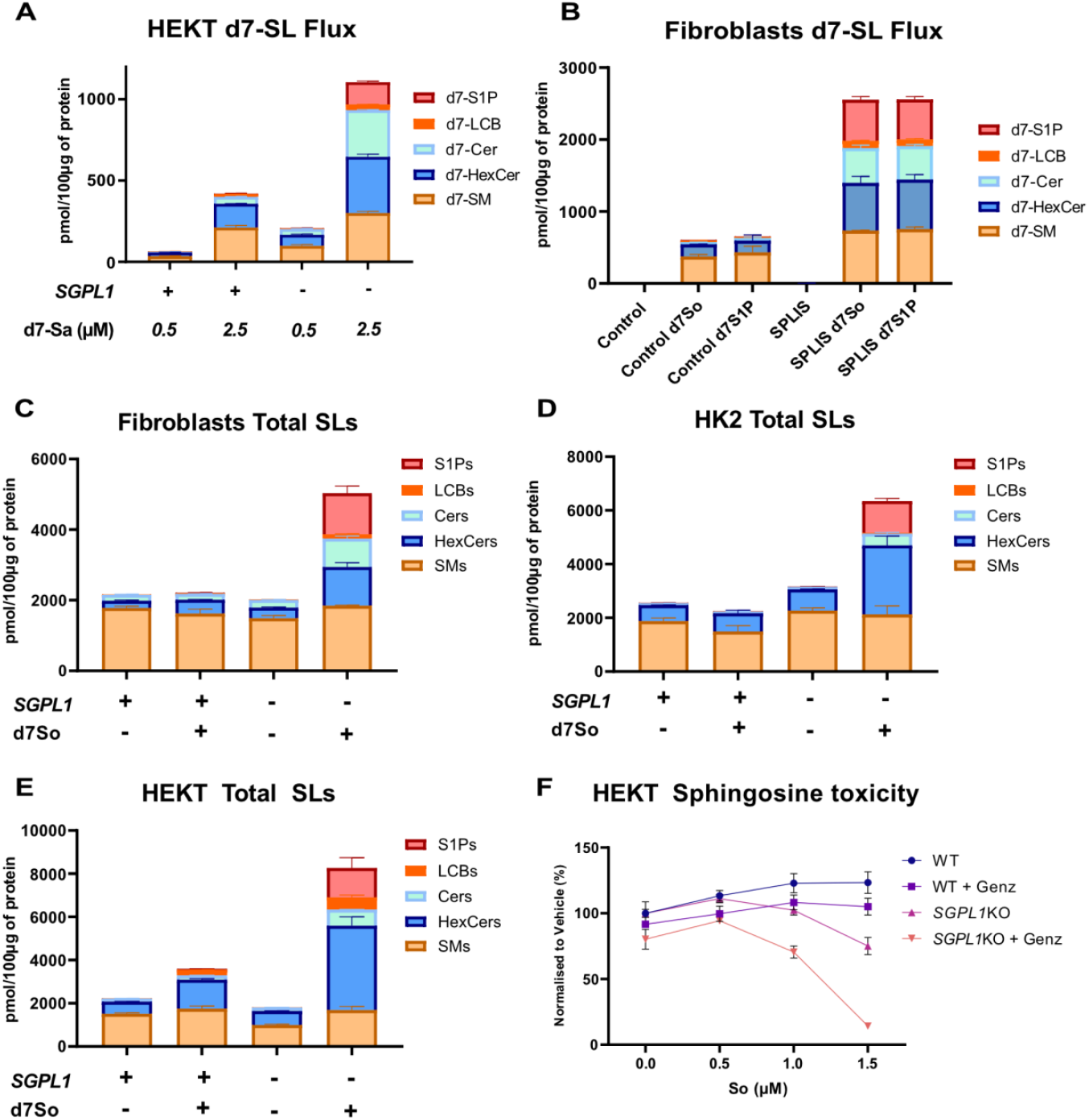
Synthesis of higher-order sphingolipids (SL) acts as an “escape” mechanism to prevent toxic S1P accumulation in SPLIS. **(A)** d7-SL profiles following supplementation with d7-sphinganine (d7-Sa, 0.5 μM or 2.5 μM) in *HEK293T* SGPL1 KO and WT cells. **(B)** d7-SL levels in primary SPLIS fibroblasts and control fibroblasts after treatment with vehicle, d7-sphingosine (d7-So, 0.5 μM), or d7-sphingosine-1-phosphate (d7-S1P, 0.5 μM) for 24 hours. **(C–E)** Total SL levels in three different SGPL1-deficient cell lines—primary fibroblasts (**C**), HK2 cells (**D**), and *HEK293T* cells (**E**)—compared to corresponding controls after 24-hour supplementation with vehicle (MeOH) or d7-So (2.0 μM). Total SL levels were calculated as the sum of d7-labeled and unlabeled SL species. Bar and stacked plots represent mean ± SD (N=3) for the indicated SL classes. Galactosylceramides and glucosylceramides are cumulatively represented as hexosylceramides (HexCer). SL levels were quantified via LC-MS/MS following lipid extraction. **(F)** ATP-based sphingosine (So) toxicity assay in *HEK293T* SGPL1 KO and WT cells. Glucosylceramide synthesis was inhibited using the glucosylceramide synthase (GCS) inhibitor Genz-123346 (Genz). Cells were exposed to increasing concentrations of So for 72 hours, and total ATP levels were quantified using the CellTiter-Glo assay. Data from the toxicity assay were normalized to the average of vehicle-treated cells (N=4).

### SPLIS-inducing variants of *SGPL1* diminished the cellular ability to clear excess S1Ps

SGPL1 converts sphingosine-1-phosphate (S1P) into hexadecenal and phosphoethanolamine. Hexadecenal is further oxidized by aldehyde dehydrogenase 3A2 (ALDH3A2) into a fatty acid, which is subsequently metabolized into other lipids, including glycerophospholipids such as phosphatidylcholine (PC) (Figure 4A) (21). Increasing concentrations of supplemented d7-sphinganine (d7-Sa) correlated with the formation of d7-labeled PCs in WT cells. This conversion was absent in SGPL1 KO cells, which instead showed a marked accumulation of d7-S1P. Additionally, other d7-labeled downstream phospholipids, such as d7-phosphatidylethanolamines (d7-PEs), d7-triglycerides (d7-TGs), and d7-diacylglycerols (d7-DGs), were detected (data not shown). Similarly, when primary SPLIS fibroblasts were supplemented with either d7-sphingosine (d7-So) or d7-S1P, WT cells efficiently converted the labeled SL into d7-PC. In contrast, SPLIS fibroblasts primarily converted the labeled SL into d7-S1P, with minimal formation of d7-PC (Figure 4C).

**Figure 4.**
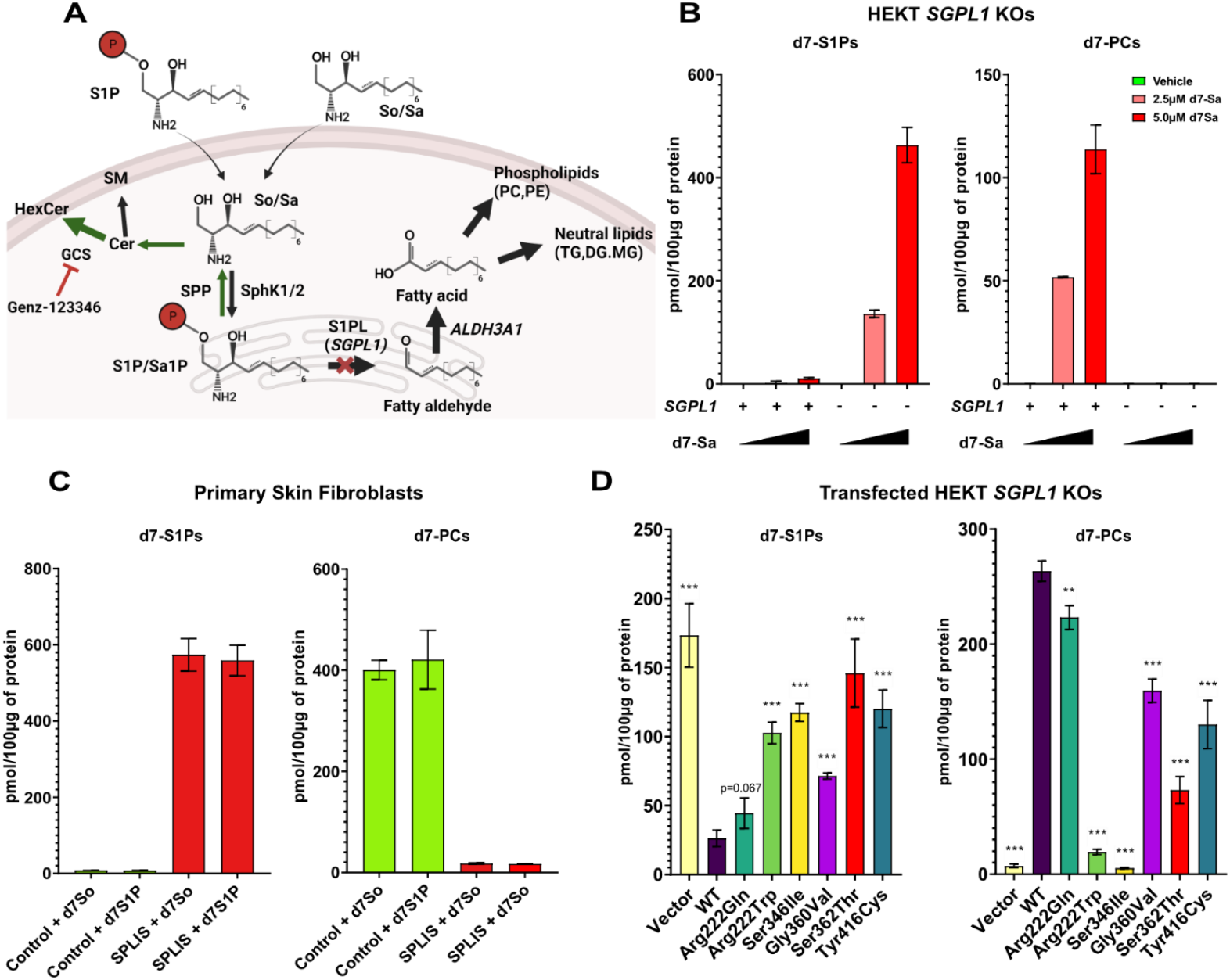
SPLIS inducing variants of *Sgpl1* diminish cell ability to clear excess S1Ps. **Figure 4**: Analysis of sphingolipid metabolism and the impact of SGPL1 mutations. **(A)** Schematic representation of the metabolism of externally supplemented sphingolipids (SL). **(B)** Levels of d7-S1P and d7-PC after 24 hours of incubation with increasing concentrations of d7-sphinganine (d7-Sa) in *HEK293T* SGPL1 KO (SGPL1 -) and WT (SGPL1 +) cells. Increasing d7-Sa concentrations correlate with higher d7-PC levels in WT cells, a capability diminished in SGPL1 KO cells, which instead show an accumulation of d7-S1P. **(C)** Levels of d7-S1P and d7-PC after 24 hours of incubation with vehicle, d7-sphingosine (d7-So, 0.5 μM), or d7-S1P (0.5 μM) in SPLIS fibroblasts compared to control fibroblasts. **(D)** Levels of d7-S1P and d7-PC after 24 hours of incubation with d7-Sa (2.0 μM) in *HEK293T* SGPL1 KO cells expressing WT SGPL1, an empty vector (EV), or six SPLIS-associated SGPL1 variants. Bar plots represent mean ± SD (N=3). d7-S1P and d7-PC levels were analyzed using LC-MS/MS following lipid extraction. Statistical significance was calculated between WT SGPL1-expressing cells and individual SGPL1 mutant/empty vector-expressing cells using a two-tailed *t*-test. ** = p < 0.005; *** = p < 0.0005.

Different SPLIS mutations are associated with varying disease severities. To investigate this, we compared the activity of six SGPL1 variants by expressing them in *HEK293T* SGPL1 KO cells (Figure 4D) and analyzing their ability to convert d7-Sa into d7-PC. All disease-associated variants demonstrated a reduced capacity to degrade d7-S1P and to form d7-PC. Notably, the most prevalent SPLIS variant, p.Arg222Gln (6), exhibited the highest residual activity (approximately 10%) and the highest protein expression (Supplementary Figure 2A, B).

### *Sgpl1* KO mice accumulate large quantities of S1P in kidneys and urine

SGPL1p deficiency results in an inability to efficiently clear exogenously supplemented sphingolipids (SL). Intracellularly, S1P levels are typically maintained in the nanomolar range, while S1P is highly abundant in plasma (∼0.5 μM). Since the kidneys process large volumes of blood, they are consistently exposed to high levels of extracellular S1P. To prevent toxic intracellular accumulation of S1P, kidney cells must actively degrade this metabolite.

*Sgpl1-/-* mice develop nephrosis characterized by hypoalbuminemia and an elevated urine albumin-to-creatinine ratio, mirroring the pathology observed in humans. We analyzed total S1P content in the kidney, liver, and muscle of WT, *Sgpl1+/-*, and *Sgpl1-/-* mice. Compared to WT animals, *Sgpl1-/-* mice showed increased S1P levels in tissues and urine (Figures 5A–D). The most pronounced S1P accumulation was observed in the kidneys of *Sgpl1-/-* mice, which is the primary organ affected in SPLIS. Glomerulosclerosis in these mice was confirmed histologically (Figure 5E). In contrast, heterozygous *Sgpl1+/-* mice did not exhibit significant S1P accumulation, suggesting that a single functional allele is sufficient to maintain normal S1P levels.

**Figure 5.**
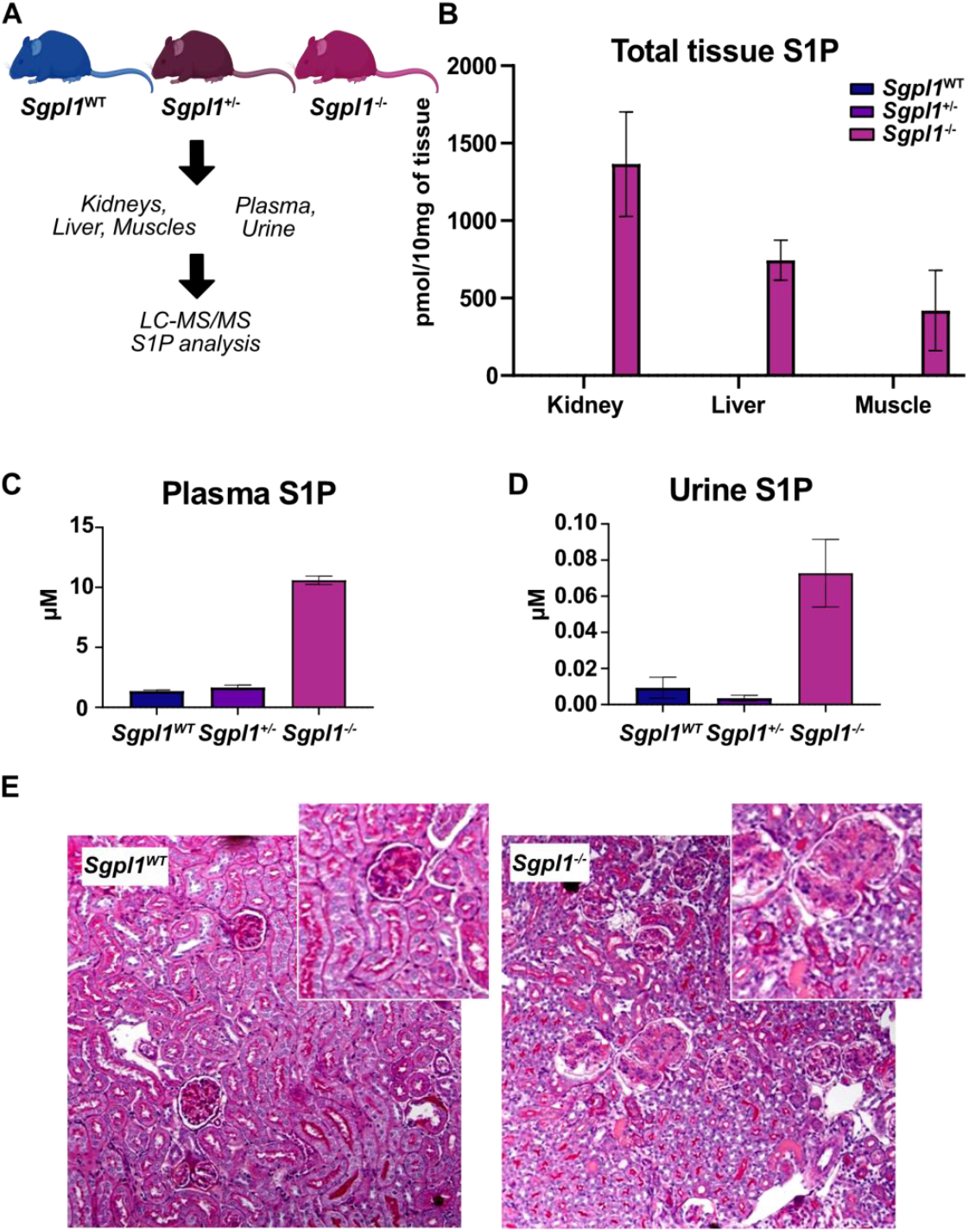
*Sgpl1* KO mice accumulate large amounts of S1P in kidneys and urine. **Figure 5**: Analysis of S1P levels and kidney pathology in *Sgpl1* mice. **(A)** Schematic representation of tissue and body fluid sample collection from *Sgpl1* WT, *Sgpl1+/-*, and *Sgpl1-/-* mice. **(B)** Total S1P levels in tissues from *Sgpl1* WT (N=3), *Sgpl1+/-* (N=3), and *Sgpl1-/-* (N=3) mice. **(C)** Total plasma S1P levels in *Sgpl1* WT (N=3), *Sgpl1+/-* (N=3), and *Sgpl1-/-* (N=3) mice. **(D)** Total urinary S1P levels in *Sgpl1* WT (N=3), *Sgpl1+/-* (N=3), and *Sgpl1-/-* (N=3) mice. Bar plots represent mean ± SD (N=3). S1P levels were measured using LC-MS/MS. Statistical significance was calculated using a two-tailed *t*-test. Differences between WT and *Sgpl1-/-* were highly significant (*p* < 0.0001). **(E)** Kidney cortex histology of wild-type (WT) and SGPL1 knockout (KO) mice stained with periodic acid-Schiff (PAS) stain. (Left, with inset detail): WT kidney cortex shows uniform glomeruli with normal size and cellularity. (Right, with inset detail): SGPL1 KO kidney cortex displays protein casts and glomeruli with heterogeneity in size and appearance, mesangial expansion, hypercellularity, and collagen deposition. Percent glomerulosclerosis: WT = 0%; KO = 37%; 50 or more glomeruli were analyzed per genotype. Scale bar = 100 μm.

The uptake of extracellular sphingoid bases is toxic to *Sgpl1*-KO cells (Figure 3F). We hypothesize that a similar mechanism underlies the specific nephrotoxic effects seen in SPLIS. Kidney cells of *Sgpl1-/-* mice absorb S1P from plasma and urine but are unable to degrade the intracellular excess of SL. The kidney contains many highly specialized cell types, some of which may be particularly vulnerable to the toxic effects of S1P due to their exposure to both excreted and reabsorbed lipids in SPLIS patients.

### Intracellular accumulation of S1P causes transient cell contraction in SPLIS fibroblasts

Exogenous supplementation with sphingosine (So) or S1P leads to intracellular S1P accumulation in SGPL1p-deficient cells (Figure 3). This increase in intracellular S1P is associated with a transient cell-rounding phenotype in SPLIS fibroblasts. Approximately 6–12 hours after the addition of So or S1P, the cells contract and subsequently re-flatten after 24 hours. This behavior was not observed in control fibroblasts (Figures 6A, B, and C).

**Figure 6.**
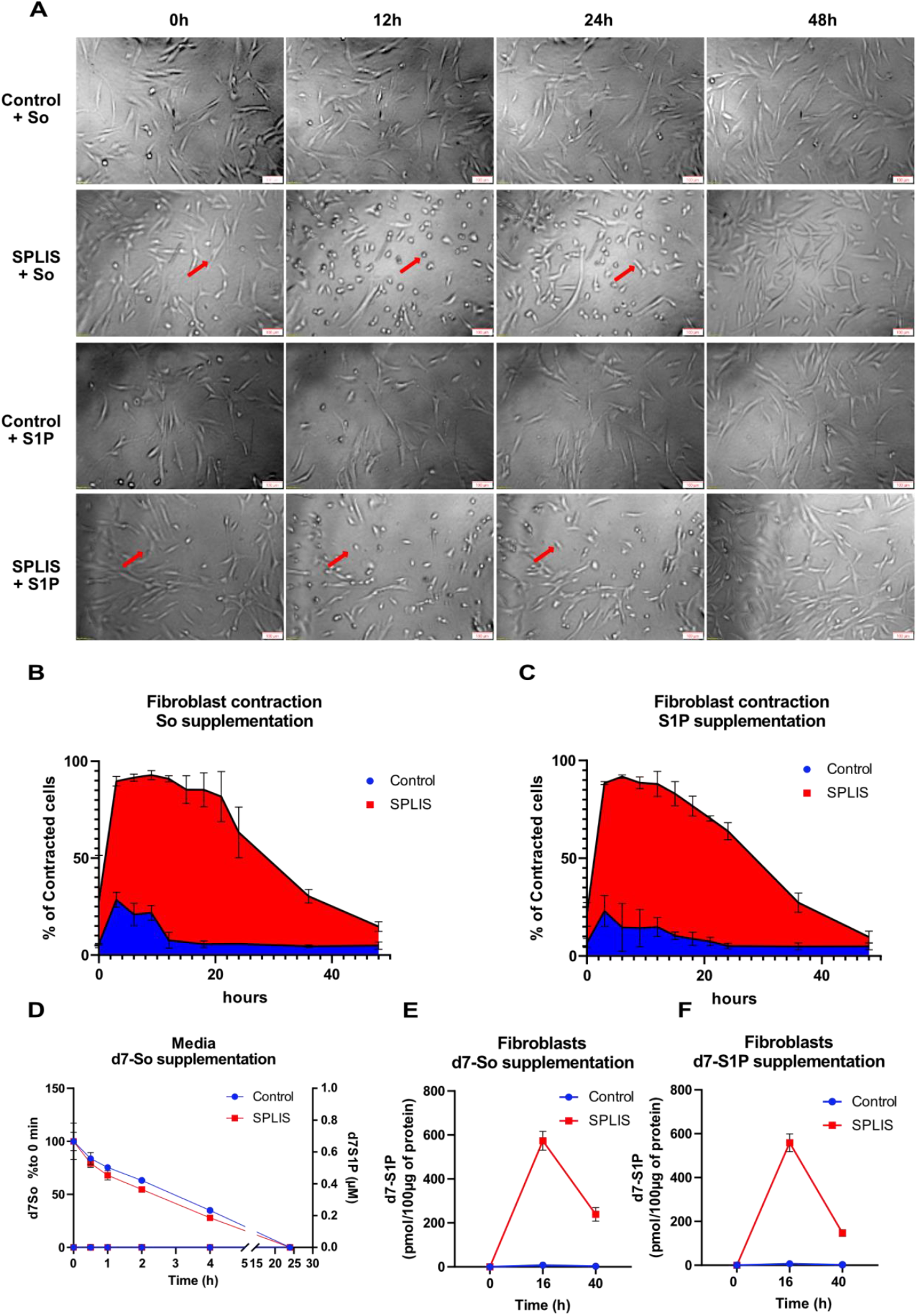
Intracellular accumulation of S1P causes cell contraction in SPLIS fibroblasts. **Figure 6**: Cytoskeletal dynamics and lipid metabolism in SPLIS fibroblasts after So or S1P supplementation. **(A)** Live-cell imaging of SPLIS fibroblasts showing cell contraction after So or S1P supplementation. Red arrows indicate contracted cells. Scale bars: 100 μm. **(B, C)** Quantification of the percentage of contracted cells per well after So or S1P supplementation in SPLIS fibroblasts and control fibroblasts. **(D)** Time-dependent uptake of d7-So (left axis) and release of d7-S1P (right axis) into the medium in cultured SPLIS and control fibroblasts. Data were normalized to the 0-hour time point. d7-So was fully absorbed within 24 hours, with no parallel release of d7-S1P into the medium. **(E)** Timeline of intracellular d7-S1P levels following supplementation with d7-So (0.5 μM) in SPLIS and control fibroblasts. **(F)** Timeline of cellular d7-S1P levels after supplementation with d7-S1P (0.5 μM) in SPLIS and control fibroblasts. Data points are represented as mean ± SD (N=3). Sphingolipid (SL) levels were analyzed using LC-MS/MS following lipid extraction.

We compared the kinetics of So and S1P absorption from the medium and the resulting intracellular S1P accumulation. Cells were incubated with d7-So or d7-S1P for 0, 16, and 40 hours. Both lipids were effectively absorbed and cleared from the medium within 24 hours at similar rates (Figure 6D). We did not detect d7-S1P release into the medium, indicating that the absorbed lipids were primarily metabolized intracellularly. In d7-So-treated SPLIS fibroblasts, intracellular d7-S1P levels increased transiently during the first 16 hours and returned to baseline by 40 hours (Figure 6E). This transient rise in intracellular S1P was closely associated with the rounding phenotype in SPLIS cells, but not in WT fibroblasts. A similar kinetic profile was observed when supplementing d7-S1P (Figure 6F).

These findings suggest that the transient rise in intracellular S1P influences the cytoskeletal dynamics of the cells, which may be relevant to the pathogenesis of SPLIS. In fact, supplementation with S1P or So in a cell migration assay confirmed the transient contraction phenotype, with re-flattening after 24 hours coinciding with the initiation of cell migration. Control fibroblasts did not show this behavior (Extended Figure 6A).

The addition of FB1 caused an increase in Sa1P levels in SPLIS fibroblasts (Figure 2E), leading to progressive cell rounding and eventual cell death, while control fibroblasts remained unaffected. This FB1-induced phenotype was rescued by co-treatment with the SPT inhibitor Myriocin, supporting the notion that the toxic response to FB1 is a consequence of de novo SL synthesis (Extended Figure 6B).

### Extended Figure 6

**Extended Figure 6..**
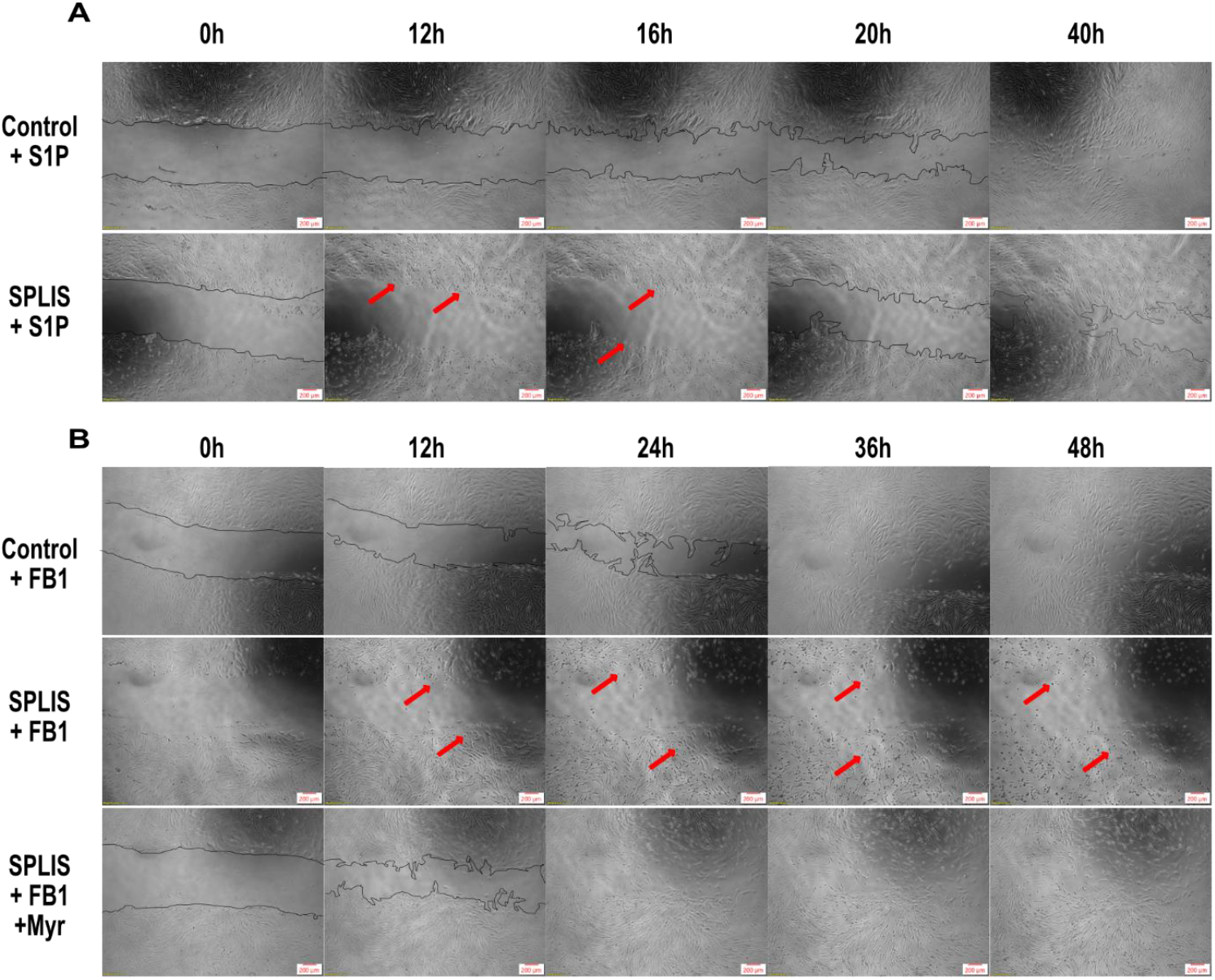
Impact of S1P and inhibitors on fibroblast migration and cytoskeletal dynamics. **(A)** Scratch assay of SPLIS primary fibroblasts and control fibroblasts supplemented with S1P (0.5 μM). **(B)** Scratch assay of SPLIS primary fibroblasts and control fibroblasts treated with the ceramide synthase inhibitor fumonisin B1 (FB1, 35 μM) or a combination of FB1 and the SPT inhibitor myriocin (Myr, 2 μM). Red arrows indicate contracted cells. Scale bars: 200 μm.

### Inhibiting ROCK prevents cytoskeletal changes in SPLIS fibroblasts and a SGPL1-deficient HK2 cell line

S1P signals through five G-protein-coupled sphingosine-1-phosphate receptors (S1PR1–5), with S1PR2 playing a crucial role in cytoskeletal regulation. Upon S1P binding, S1PR2 activates the Rho-associated coiled-coil-containing protein kinase (ROCK) pathway, which promotes actin stress fiber assembly and focal adhesion formation. These processes are essential for maintaining cell shape, motility, and stability. To determine whether the observed cytoskeletal effects of S1P in SPLIS fibroblasts are linked to this pathway, we inhibited ROCK using the specific inhibitor Fasudil. Additionally, we compared the effects of Fingolimod (FTY720), an antagonist of S1PR1, 3, 4, and 5 (but not S1PR2), and JTE013, a specific S1PR2 antagonist (22). The contraction phenotype was assessed in an SGPL1-deficient human proximal tubule cell line (HK2) using phalloidin staining (Figure 7). Inhibiting the Rho-ROCK axis with Fasudil prevented the contraction phenotype in SPLIS cells. JTE013 showed a partial rescue effect, while FTY720 had no effect, indicating that S1PR2 is responsible for the contraction phenotype. For quantitative analysis, we used the software tool CellProfiler. Cell contraction was quantified per cell and defined by the function 1/log_10_(Cell surface/Nucleus surface). Averages from four images per well were used for the analysis. SPLIS fibroblasts exhibited significantly increased cell rounding in response to S1P treatment, which was significantly reduced in the presence of Fasudil (Figure 7B). Finally, we tested the effect of Fasudil on HK2 kidney cells. SGPL1 KO or WT HK2 cells were cultured with increasing concentrations of S1P in the presence or absence of Fasudil. In SGPL1 KO HK2 cells, S1P disrupted the formation of a coherent epithelial layer, a defect that was reversed when Fasudil was co-administered (Figure 7C). In summary, these findings suggest that inhibition of the Rho-ROCK signaling axis can rescue SPLIS-induced cytoskeletal phenotypes.

**Figure 7.**
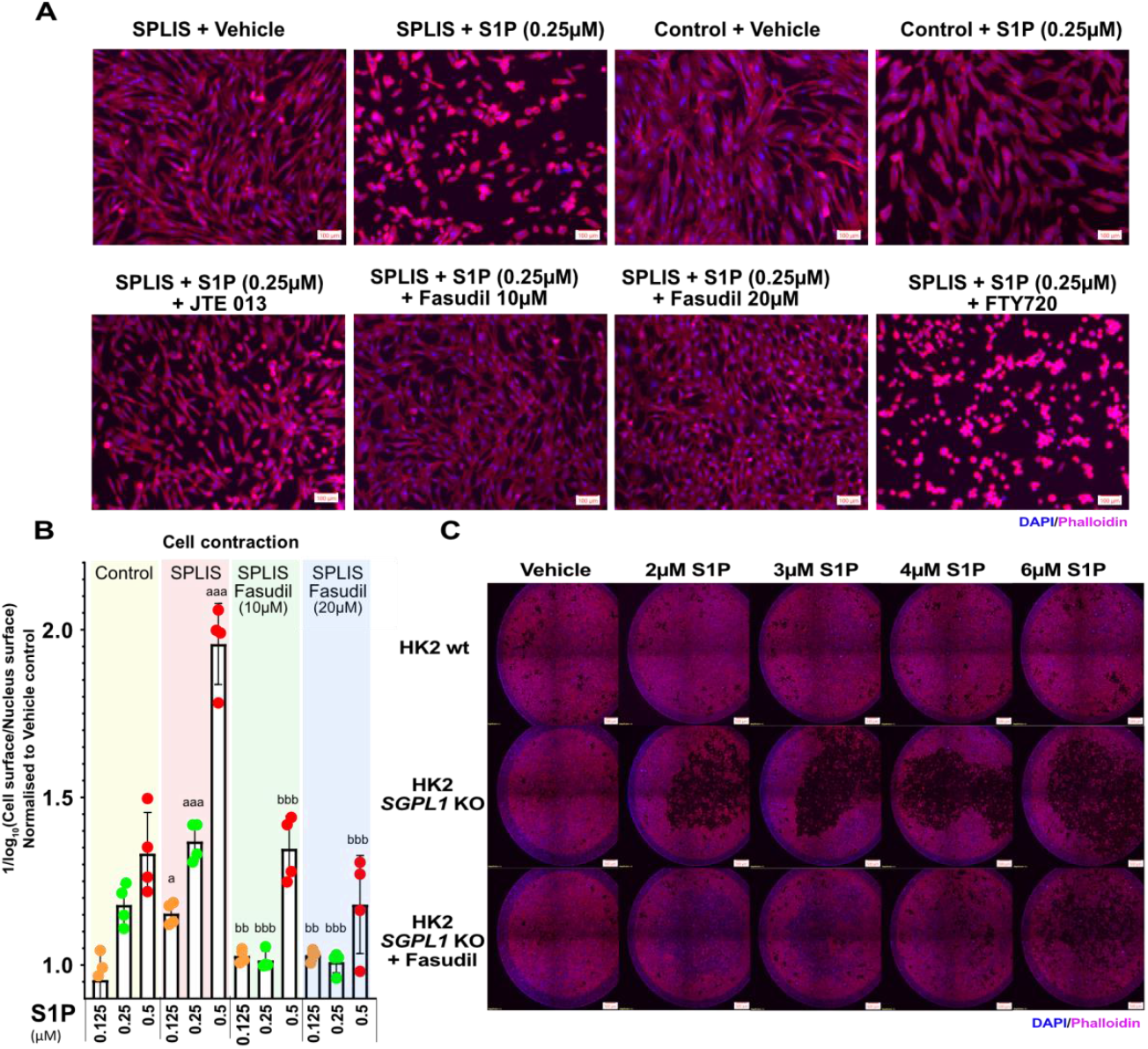
Fasudil rescues S1P induced cytoskeletal phenotypes in SPLIS fibroblasts and SGPL1-deficient HK2 cell line. **Figure 7**: Cytoskeletal and epithelial effects of S1P and ROCK inhibition in SPLIS fibroblasts and HK2 cells. **(A)** Fluorescence imaging of SPLIS primary fibroblasts and control fibroblasts treated with vehicle (MeOH) or S1P (0.25 μM) for 6 hours. Cells were also treated with the ROCK inhibitor (Fasudil), the sphingosine-1-phosphate receptor 2 inhibitor (JTE013), or the sphingosine-1-phosphate receptor 1 modulator (Fingolimod, FTY720). After treatment, cells were fixed with 4% PFA and stained with Phalloidin (actin) and DAPI (nucleus). Scale bars: 100 μm. **(B)** Quantification of cell contraction. Compiled images of whole wells were analyzed using CellProfiler software. Cell contraction was defined by the formula 1/log10(Cell surface/Nucleus surface). Values were normalized to the average of vehicle-treated cells. Data are represented as mean ± SD (N=4). Statistical analysis was performed using a two-tailed parametric t-test. a: Comparison between control and SPLIS fibroblasts at the same S1P concentration (a = p < 0.05, aa = p < 0.005, aaa = p < 0.0005). b: Comparison between SPLIS fibroblasts treated without or with two concentrations of Fasudil (ROCK inhibitor) (b = p < 0.05, bb = p < 0.005, bbb = p < 0.0005). **(C)**Impairment of renal epithelium formation in SGPL1 KO human proximal tubule cells (HK2) after S1P supplementation. SGPL1 KO or WT HK2 cells were grown for 72 hours in the presence of increasing S1P concentrations, with or without Fasudil, as indicated. Cells were fixed with 4% PFA and stained with Phalloidin (actin) and DAPI (nucleus). Whole wells were imaged using fluorescence microscopy. Representative images from three independent replicates are shown. Scale bars: 500 μm.

## Discussion

SPLIS, or nephrotic syndrome type 14, is an inherited disease caused by recessive mutations in *SGPL1*. Since SGPL1p is responsible for the terminal degradation of sphingolipids (SL), reduced SGPL1p activity is expected to result in increased intracellular SL levels. However, the underlying metabolic rearrangements caused by SGPL1 insufficiency remain poorly understood. Additionally, the pathophysiological mechanisms of SPLIS leading to nephrosis are not well characterized, and no targeted therapies have been proposed. Here, we report the case of a patient with a novel SPLIS-associated *SGPL1* variant (*p*.*Ser362Thr*). The patient presented with congenital nephrotic syndrome and immunodeficiency, resulting in early death at 9 months of age. Contrary to previous studies (23), we did not observe significantly increased S1P levels in the patient’s plasma and found only slightly elevated intracellular S1P levels in patient-derived skin fibroblasts. Instead, we detected significantly increased levels of ceramide (Cer), sphingomyelin (SM), and hexosylceramide (HexCer).

Sphingolipid (SL) de novo formation is initiated by serine palmitoyltransferase (SPT) and regulated by its subunits ORMDL1-3 in response to intracellular ceramide levels (18, 24). Increasing intracellular SL content by supplementing sphingosine, sphinganine, or membrane-permeable C6-ceramide reduced SPT activity and lowered SL de novo synthesis in both WT and SGPL1p-deficient cells (20). In fact, SPT activity and SL de novo synthesis are significantly reduced in SGPL1p-deficient cells, including patient-derived fibroblasts. Reduced SPT activity has also been reported in SGPL1p-deficient neurons (25). We hypothesize that in SPLIS, the impaired breakdown of SL is partially offset by a decrease in SL *de-novo* synthesis, thereby helping to maintain low intracellular S1P levels despite the absence of SGPL1p activity. In contrast, inhibiting ceramide synthesis with FB1 or lowering ORMDL1-3 expression, significantly increases SPT activity. Under conditions of increased SL synthesis, SGPL1p-deficient cells accumulated significant amounts of S1P, which was not observed in cells expressing functional SGPL1p. Conversely, S1P accumulation was prevented when C6-ceramide was supplemented in addition to FB1. These findings suggest that in the absence of SGPL1p, SPT regulation via the ceramide-ORMDL axis is crucial for maintaining baseline SL levels. This hypothesis is supported by data from *Drosophila*, where the muscle phenotypes in flies with hypomorphic mutations in *Sply* (the fly orthologue of SGPL1) were attenuated by overexpressing *Lace* (the orthologue of SPT) (26). The importance of a controlling SPT and SL-*de novo* synthesis in absence of SGPL1p activity is further supported by the higher toxicity of FB1 in SGPL1p-deficient cells (Figure 2G). Typically, we did not observe a significant increase in intracellular S1P or a notable release of S1P into the medium in the absence of SGPL1p (Figure 6D). However, we observed a significant increase in hexosylceramide (HexCer), suggesting that the increased synthesis of glycosphingolipids may serve as a secondary compensatory mechanism to prevent intracellular accumulation of S1P and other potentially harmful SL species. This protective mechanism might explain why supplementing low amounts of d7-Sa to SGPL1p-deficient cells increased HexCer levels without altering S1P, while higher concentrations of d7-Sa overloaded the system’s compensatory capacity, resulting in a toxic increase in S1P. A compensatory conversion of ceramide into HexCer has also been reported in SGPL1p-deficient neurons (25). SGPL1p-deficient cells supplemented with external SL exhibited the highest HexCer levels. However, blocking HexCer synthesis with the glucosylceramide synthase (GCS) inhibitor *Genz-123346* sensitized SGPL1p-deficient cells to sphingosine supplementation, further supporting the role of HexCer synthesis as a compensatory mechanism.

SGPL1p converts S1P into fatty aldehydes, which are subsequently metabolized into fatty acids and ultimately incorporated into glycerophospholipids (e.g. PCs PEs) and neutral lipids (e.g. TGs and DGs) (21). In SGPL1-deficient cells, the inability to efficiently cleave S1P, results in reduced conversion into glycerophospholipids, particularly when cells are supplemented with external LCBs such as d7-So, d7-Sa or d7-S1P (Figure 4). This impaired conversion has been linked to reduced PE formation in SPLIS, which may compromise autophagy, as lipidation of LC3—a key step in autophagosome formation—relies on adequate PE levels (27).

The SPLIS variants of SGPL1p reduce enzyme activity (Figure 4D) which affects the conversion of externally supplemented SL into PCs (Figure 4D). The highest residual activity was observed for SGPL1 p.Arg222Gln, whic is the most frequently reported SPLIS variant and is associated with a comparatively mild phenotype (7) (6).

We observed significant accumulation of S1P in the tissues, blood, and urine of *Sgpl1-/-* mice. The most pronounced S1P accumulation was in the kidneys, the primary organ affected in SPLIS. Kidneys are constantly exposed to relatively high S1P concentrations in the blood (approximately 0.5 μM), creating persistent pressure to degrade the reabsorbed S1P. Elevated S1P levels in the urine of *Sgpl1-/-* mice suggest that in SPLIS, tubular kidney cells exposed to high extracellular S1P levels, but incapable of degrading it, may be particularly vulnerable to its toxic effects. Given that S1P reabsorption from urine is specific to the kidneys, this inability to degrade S1P likely explains the selective nephrotic phenotype observed in SPLIS.

The supplementation of external sphingolipids (So or S1P) to SPLIS primary fibroblasts caused profound cytoskeletal changes, manifesting as a time-dependent but transient cell contraction closely correlated with changes in intracellular S1P levels. Similarly, supplementation of S1P induced epithelial defects in SGPL1p-deficient HK2 cells. S1P is a pharmacologically potent molecule that exerts its effects through five specific receptors (S1PR1–5) (28). S1PR2 links to cytoskeletal regulation via the Rho/ROCK signaling pathway (29) and altered S1PR2 signaling has been linked to kidney disease previously (30). Blocking S1PR2 with JTE013 and inhibiting the ROCK signaling axis with Fasudil prevented S1P-induced cytoskeletal changes in SPLIS primary fibroblasts and *HK2* SGPL1 KO cells.

Fasudil is clinically approved in China and Japan for treating cerebral vasospasm and ischemia associated with subarachnoid hemorrhage. Our data suggest that Fasudil could be a potential therapeutic option for patients with SPLIS. Additionally, pharmacologic inhibition of SPT can effectively suppress SL biosynthesis and may hold therapeutic potential. The only FDA-approved SPT inhibitor, L-cycloserine, also inhibits SPL, limiting its safe use to patients with non-functional SPL (complete loss-of-function variants). However, it may be feasible to develop SPT inhibitors that specifically target SPT without affecting other B6-dependent enzymes, including SPL, and that exhibit less toxicity than myriocin.

However, substrate depletion strategies for reducing SL biosynthesis pose challenges, as lowering serine levels increases alanine utilization, which promotes the formation of neurotoxic 1-deoxy-sphingolipids. Moreover, dietary sources of sphingolipids, which are present in nearly all foods, further complicate this approach. Additional preclinical studies are needed to confirm the potential therapeutic benefits of these strategies.

---

**Supplementary Figure 1.**
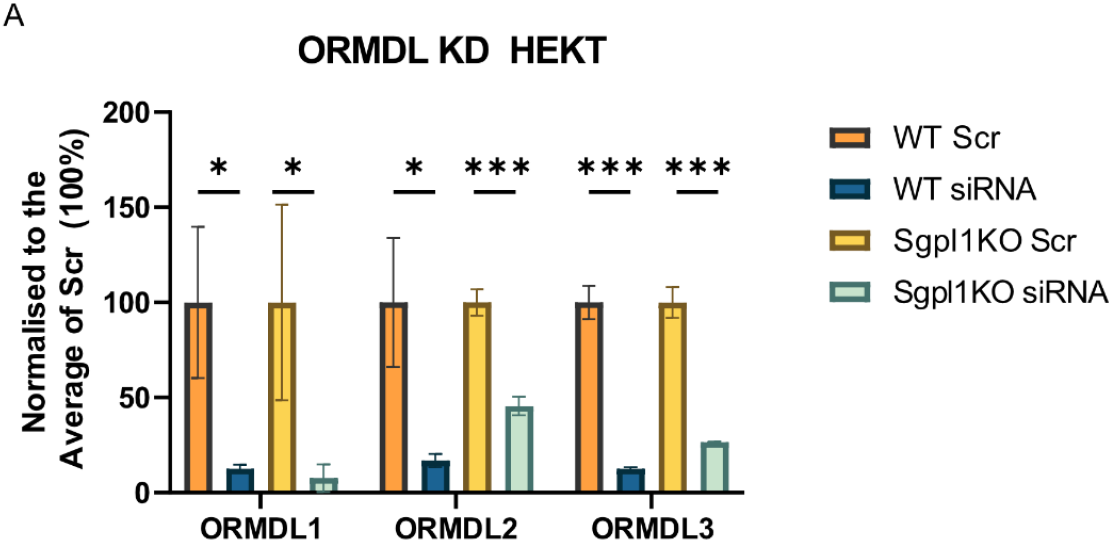
**A** RMDL expression after knockdown (KD) with ORMDL 1, 2, 3, siRNA or non-coding Scramble (Scr) as quantified by qRT-PCR. Total RNA levels were calculated from a dilution curve and normalized to GAPDH. Levels were normalized to Scr. In total 3 tests were performed. Multiple testing was corrected using two-stage step-up method (Benjamini, Krieger and Yekutieli). *=q<0.05, **=q<0.005, ***=q<0.0005.

**Supplementary Figure 2.**
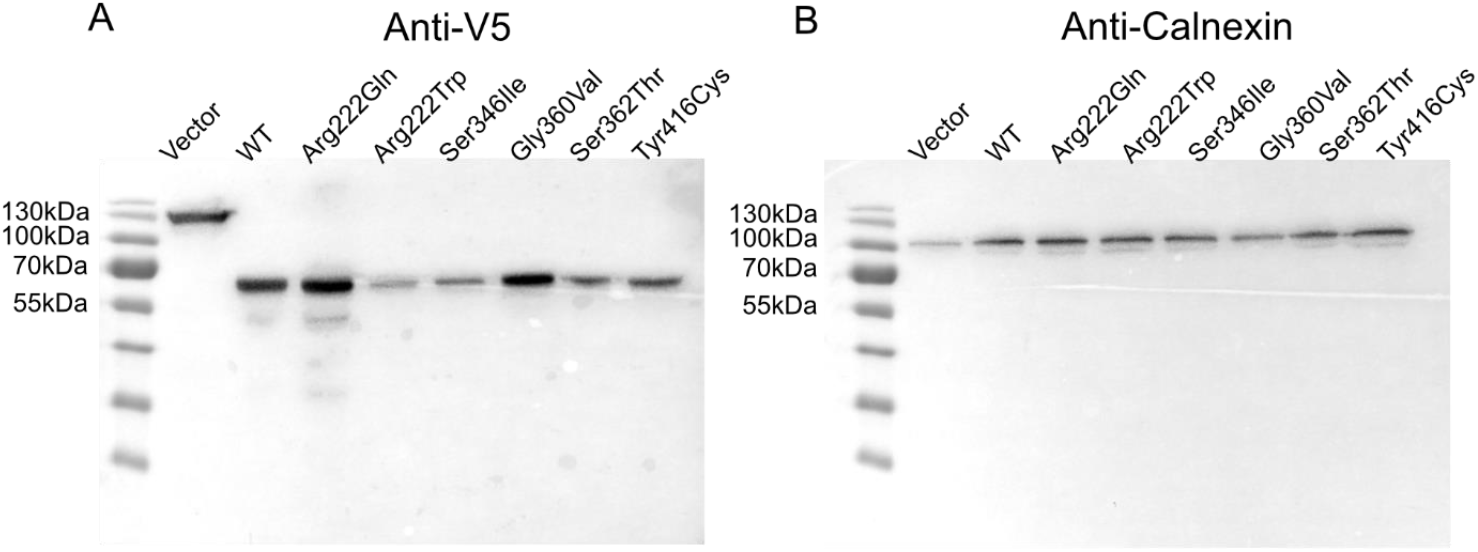
A P.Lenti-Sgpl1-V-5 expression of different SPLIS inducing SGPL1 mutants in HEKT SGPL1 KO background. P.Lenti-beta-Galactosidase-V5 expressed in HEKT SGPL1 KO was used as Vector control. **B** Anti-Calnexin expression as a loading control

